# Population-based Carrier Screening and Prenatal Diagnosis of Fragile X Syndrome in East Asian Populations

**DOI:** 10.1101/2020.09.10.292219

**Authors:** Qiwei Guo, Yih-Yuan Chang, Chien-Hao Huang, Yu-Shan Hsiao, Yu-Chiao Hsiao, I-fan Chiu, Yulin Zhou, Haixia Zhang, Tsang-Ming Ko

## Abstract

Identification of carriers of fragile X syndrome (FXS) with the subsequent prenatal diagnosis, and knowledge of FXS-associated genetic profiles are essential for intervention in specific populations. We report the results of carrier screening of 39,458 East Asian adult women and prenatal diagnosis from 87 FXS carriers. The prevalence of FXS carriers and incidence of full mutation fetuses in carrier pregnancies were found to be 1/556 and 11.0%, respectively. The prevalence of FXS carriers and full mutation fetuses was estimated to be 1/581 and 1/3124 in East Asian populations, respectively. We confirmed the validity of the current threshold of CGG repeats for *FMR1* categorization; the integral risks of full mutation expansion were approximately 6.0%, 43.8%, and 100% for premutation alleles with 55-74, 75-89, and ≥90 CGG repeats, respectively. The protective effect of AGG interruption in East Asian populations was validated, which is important in protecting premutation alleles with 75-89 CGG repeats from full mutation expansion. Lastly, family history was shown not an effective indicator for FXS carrier screening in East Asian populations and population-based screening was more cost-effective. This study provides an insight into the largest carrier screening and prenatal diagnosis for FXS in East Asian populations to date. The FXS-associated genetic profiles of East Asian populations are delineated and population-based carrier screening is shown to be promising for FXS intervention.

## Introduction

Fragile X syndrome (FXS) (#MIM300624) is the leading monogenic cause of autism and autism spectrum disorders and the second most common cause of inherited intellectual disability ^1^. FXS affects approximately 1 in 4,000 males and 1 in 7,000 females worldwide ^2^. The genetic basis of FXS is the translational silencing of FMRP translational regulator 1 (*FMR1*), an X-linked dominant gene with a reduced penetrance of 80% in males and 30-50% in females ^1^. *FMR1* spans 17 exons and encodes FMRP, which plays a fundamental role in the synapses and the normal development of dendrites ^3^. A segment of CGG trinucleotide repeat is located in the untranslated region of the first exon of *FMR1* and the number of CGG repeats is polymorphic ^4^. According to the guidelines of American College of Medical Genetics and Genomics (ACMG), *FMR1* is categorized into four allelic forms based on the number of CGG repeat: normal, intermediate, premutation (PM), and full mutation (FM) ^5^. Specifically, normal alleles have a range of ∼5 to ∼44 repeats and are meiotic or mitotic stable. Intermediate alleles range from ∼45 to ∼54 repeats and their repeat number can slightly increase or decrease in intergenerational transmission. PM alleles range from ∼55 to ∼200 repeats and can expand to FM during transmission from parents (almost solely from the mother) to the child. In addition, the PM allele can cause fragile X-associated tremor ataxia syndrome, which mainly affects males, and fragile X-associated primary ovarian insufficiency, which results in reduced fertility in females. FM alleles have more than 200 repeats. The adjacent CpG islands and the repeats themselves are methylated, which block the subsequent transcription and translation, and accounts for most cases of FXS. Therefore, asymptomatic women with PM or FM alleles are considered to be FXS carriers since their children are at risk of FXS. In addition to the CGG repeat number, other factors may be associated with the risk of CGG intergenerational expansion, although their effects require further confirmation ^6-9^. Among these, AGG interruption, which refers to AGG triplet within the CGG repeats, has received the most attention ^6, 10, 11^. In many cases, CGG repeats are interrupted by one or more AGG interruptions. AGG interruption has been suggested to act as a protective factor, anchoring the region during replication to prevent strand slippage, thus decreasing the risk for intergenerational CGG expansion ^12^. Notably, genetic profiles, such as the prevalence of carrier and FXS, the intergenerational transmission profile of CGG repeat, and the protective effect of AGG interruption, vary widely among different populations ^5, 11, 13, 14^.

Considering the significant morbidity associated with FXS and the absence of an effective treatment, the prenatal diagnosis and intervention of FXS is warranted ^5^. In this context, the identification of FXS carriers is an essential work for FXS intervention ^15^. Moreover, evaluating the FXS-associated genetic profiles for specific populations is also fundamental to improve our knowledge for genetic counselling, revising the *FMR1* categorization, refining risk assessment, and policy-making for carrier screening strategies.

In this study, we reported the results of FXS genetic testing over a period of six years. Our analysis delineated the FXS-associated genetic profiles in East Asian populations. Furthermore, we compared the cost-effectiveness of FXS carrier screening strategies in general or targeted populations.

## Materials and Methods

### Subjects

We retrospectively analyzed the results of FXS genetic testing from January 1, 2014 to December 31, 2019 in Ko’s Obstetrics & Gynecology Clinic, Taiwan. We enrolled 3 cohorts of subjects in this study. Cohort 1 included 39,458 East Asian adult women receiving testing for the first time, regardless of whether they had a family history of FXS-associated conditions. Cohort 2 included 34 East Asian adult women who had been diagnosed as FXS carriers in other institutes and were seeking prenatal diagnosis in our clinic. Cohort 3 comprised the prenatal diagnosis cases in Cohorts 1 and 2.

### FXS genetic testing

Pre- and pos-test counseling was offered to every participant by well-trained genetic counsellors, according to the ACMG Standards and Guidelines. The CGG repeat and AGG interruption status of the *FMR1* was determined using AmplideX^®^ FMR1 PCR reagents (Asuragen, Austin, TX) with capillary electrophoresis on ABI3730xl instruments (Thermo-Fisher Scientific, Waltham, MA). To ensure the accuracy of this study, all PM or FM samples were re-tested and confirmed using Biofast^®^ FMR1 PCR reagents (Bioson, Xiamen, China).

### Reference data

Data from previous studies involving detection of FXS in East Asian populations were combined and analyzed to estimate the FXS-associated genetic profiles.

### Samples

For each adult woman, 2 mL of EDTA-anticoagulated peripheral blood was sampled.

For each prenatal diagnosis, approximately 10 mg of villus or 10 mL of amniotic fluid was sampled. DNA was extracted using the MagCore Automatic Nucleic Acid Extractor (RBC Bioscience, Taipei, Taiwan), according to the manufacturer’s protocols. The DNA concentration was determined by measuring the absorbance at 260 nm using a NanoDrop 1000 spectrophotometer (Thermo Fisher, Waltham, MA)

### Cost-effectiveness analysis

We developed a simplified decision-analytic model to compare the cost-effectiveness of the FXS carrier screening strategies for adult women in cohort 1 that was based on different subjects: all adult women (population-based screening), and high-risk women with a family history of fragile X-related disorders or intellectual disability suggestive of FXS (targeted screening) ^15^. In this model, we assumed that the accuracy of FXS genetic testing was 100% with a penetrance of *FMR1* of 60% (average of male and female) ^1^. The cost of FXS carrier screening in Taiwan is approximately 100 USD per case (including genetic testing and counseling), and the cost of prenatal diagnosis is approximately 400 USD per case (including sampling, FXS genetic testing, and maternal cell contamination identification). Since no data regarding the expenses of FXS treatment is available in Taiwan, we conservatively assumed that the lifecycle cost for an FXS patient would be approximately 500,000 USD, based on data from the Europe and the United States ^16, 17^.

In our cost-effectiveness analysis, the cost is denoted as the sum of the estimated cost for carrier screening and prenatal diagnosis. The potential cost is the estimated lifecycle cost for FXS patients that would be born in a targeted screening branch. The effectiveness of a strategy was calculated by dividing the total number of identified FM fetuses by the total number of FM fetuses that would be generated by adult women in cohort 1.

### Statistical analyses

Normality testing was performed to determine whether a dataset followed a normal distribution. When a data set was non-normally distributed, a non-parametric method, Kruskal–Wallis analysis of variance (ANOVA), was used to test whether samples originated from the same distribution using OriginPro software (version 8.0, OriginLab Corp., Northampton, MA). The Chi-square test was performed using SPSS statistical software (version 20.0; IBM, Armonk, NY) to determine the differences for the frequencies between or among the different datasets.

### Ethics statement

Signed informed consent was obtained from each participant. All participants agreed to the use of their data for research purposes. Except for the results of genetic testing and fetal genders, all information, including the names of the participants, were de-identified. This study was approved by the Research Ethics Committees of the Women and Children’s Hospital at the School of Medicine of Xiamen University and the Ko’s Obstetrics & Gynecology Clinic.

## Results

### Overview of FXS genetic testing results

An overview of the FXS genetic testing results is provided in Table 1. Cohort 1 indicates the scenario of population-based FXS carrier screening. In this cohort, we identified 71 PM carriers from 39,458 adult women, revealing a 1/556 prevalence of FXS carriers in the East Asian female population. Combined with data from previous studies with relatively larger sample sizes (e.g. over 5,000) ^18-21^, the estimated prevalence of FXS carriers was approximately 1/581 and 1/796 in the general and high-risk East Asian population, respectively (Tables 2 and 3).

**Table 1.** Overview of the FXS genetic testing results.

|  | Cohort 1 | Cohort 2 | Cohort 3 |
| --- | --- | --- | --- |
| **Population-based carrier screening** |  |  |  |
| Adult women | 39458 |  |  |
| FXS carriers | 71 |  |  |
| PM carriers | 68 |  |  |
| FM carriers | 2 |  |  |
| Mosaic carrier | 1 |  |  |
| Prevalence of FXS carriers | 1/556 |  |  |
| **Prenatal diagnosis** |  |  |  |
| FXS carriers | 53 | 34 | 87 |
| Fetuses (male: female) | 57 (25:32) | 34 (20:14) | 91 (45:46) |
| Fetal *FMRI* genotypes |  |  |  |
| Normal (male: female) | 25 (9:16) | 14 (7:7) | 39 (16:23) |
| PM (male: female) | 27 (15:12) | 15 (12:3) | 42 (27:15) |
| FM (male: female) | 5 (1:4) | 5 (1:4) | 10 (2:8) |
| Incidence of FM fetuses | 8.8% | 14.7% | 11.0% |

FXS: fragile X syndrome; PM: premutation; FM: full mutation.

**Table 2.**
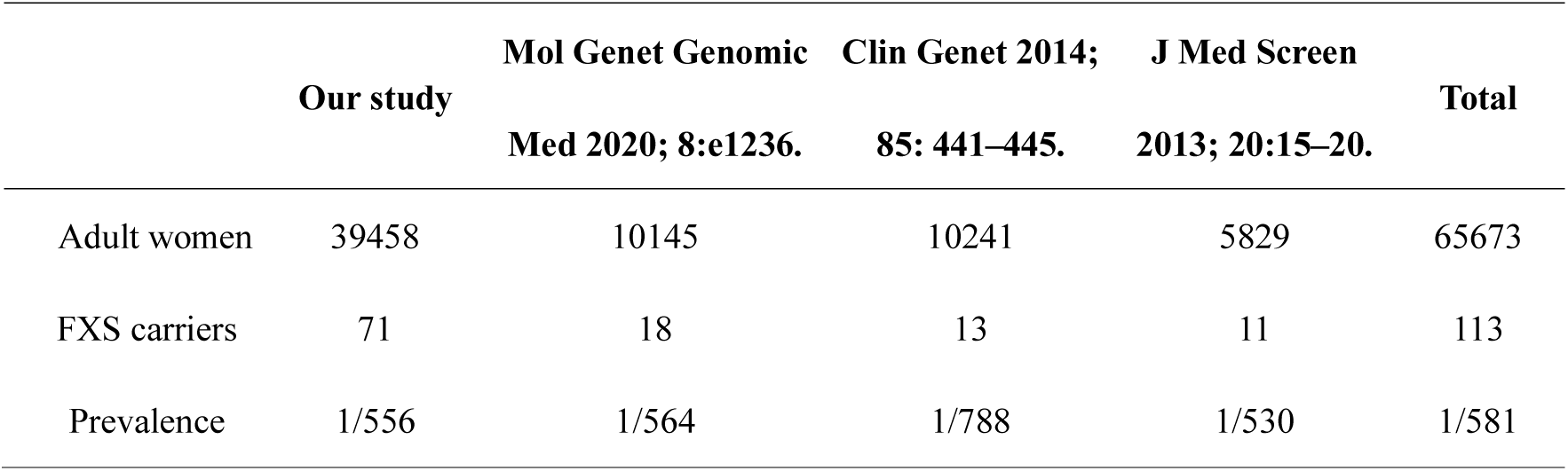
Prevalence of FXS carriers in general East Asian populations.

|  | **Our study** | **Mol Genet Genomic**<br>**Med 2020; 8:e1236.** | **Clin Genet 2014;**<br>**85: 441–445.** | **J Med Screen**<br>**2013; 20:15–20.** | **Total** |
| --- | --- | --- | --- | --- | --- |
| Adult women | 39458 | 10145 | 10241 | 5829 | 65673 |
| FXS carriers | 71 | 18 | 13 | 11 | 113 |
| Prevalence | 1/556 | 1/564 | 1/788 | 1/530 | 1/581 |

**Table 3.** Prevalence of FXS carriers in East Asian populations without FXS-associated family history.

|  | **Sci Rep 2019;**<br>**9(1): 15456.** | **Clin Genet 2014;**<br>**85: 441–445.** | **J Med Screen**<br>**2013; 20:15–20.** | **Total** |
| --- | --- | --- | --- | --- |
| Adult women | 20188 | 10146 | 5470 | 35804 |
| FXS carriers | 26 | 12 | 7 | 45 |
| Prevalence | 1/776 | 1/846 | 1/781 | 1/796 |

After genetic counselling, 53 out of 71 carriers in cohort 1 and 34 carriers in cohort 2 received prenatal diagnosis in our clinic. Combining the two cohorts (i.e. in cohort 3), 91 fetal *FMR1* genotypes were obtained from 87 carriers, since three carriers had two pregnancies and one carrier had a twin pregnancy (Supplemental Tables 1-3). The incidence of FM fetuses was 11.0% in all carrier pregnancies (Table 1). Compared to the carrier prevalence, the incidence of FM fetuses was more widely variable (9.1%-33.3%) among different studies ^19-23^. The average incidence was 18.6% (Table 4). Therefore, combining the 1/581 carrier prevalence and 18.6% incidence of the FM fetuses in carrier pregnancies, the estimated prevalence of FM fetuses in the East Asian population would be approximately 1/3124.

**Table 4.** Prenatal diagnosis for FXS carriers in East Asian populations.

|  | **Our study** | **Chin J Med Genet** | **Sci Rep 2019;** | **Clin Genet 2017;** | **Clin Genet 2014;** | **J Med Screen** | **Total** |
| --- | --- | --- | --- | --- | --- | --- | --- |
|  |  | **2019; 36(9):866-869** | **9(1): 15456.** | **92:217–220.** | **85: 441–445.** | **2013; 20:15–20.** |  |
| FXS carriers | 87 | 29 | 27 | 24 | 26 | 10 | 203 |
| Fetuses (male: female) | 91 (45:46) | 30 | 31<sup>a</sup> (15:10) | 32 | 26 | 11 | 221 |
| Fetal *FMRI* genotypes |  |  |  |  |  |  |  |
| WT (male: female) | 39 (16:23) | 17 | 13 (8:5) | 18 | 13 | 5 | 105 |
| PM (male: female) | 42 (27:15) | 3 | 11 (4:7) | 6 | 8 | 5 | 75 |
| FM (male: female) | 10 (2:8) | 10 | 7 (2:5) | 8 | 5 | 1 | 41 |
| Incidence of FM fetuses | 11.0% | 33.3% | 22.6% | 25.0% | 19.2% | 9.1% | 18.6% |

<sup>a</sup>A pregnancy in which a fetus carries a *FMRI* deletion of 5' UTR and exon 1 was excluded in this cohort.

Noticeably, unlike the gender distribution of normal and PM fetuses, female was the predominant gender in FM fetuses (Table 1).

### Intergenerational CGG repeat transmission profiles

In cohort 3 (Table 1), we identified 42 PM and 10 FM fetuses that had inherited maternal PM or FM alleles. Of these 52 transmission events, 10 (19.2%) were stably transmitted, while 42 (80.8%) increased in size (Supplemental Tables 2 and 3).

As shown in Fig. 1A, the extent of unstable transmissions and FM expansions increased markedly with an increasing maternal CGG repeat size. The smallest size of PM allele associated with FM expansion was 75, while all PM alleles ≥90 CGG repeats expanded to FM alleles. Combined with previous studies ^19-21, 23^, the integral risk of FM expansion was determined to be 6.0%, 43.8%, and 100% for PM alleles with 55-74, 75-89, and ≥ 90 CGG repeats, respectively (Table 5).

**Table 5.**
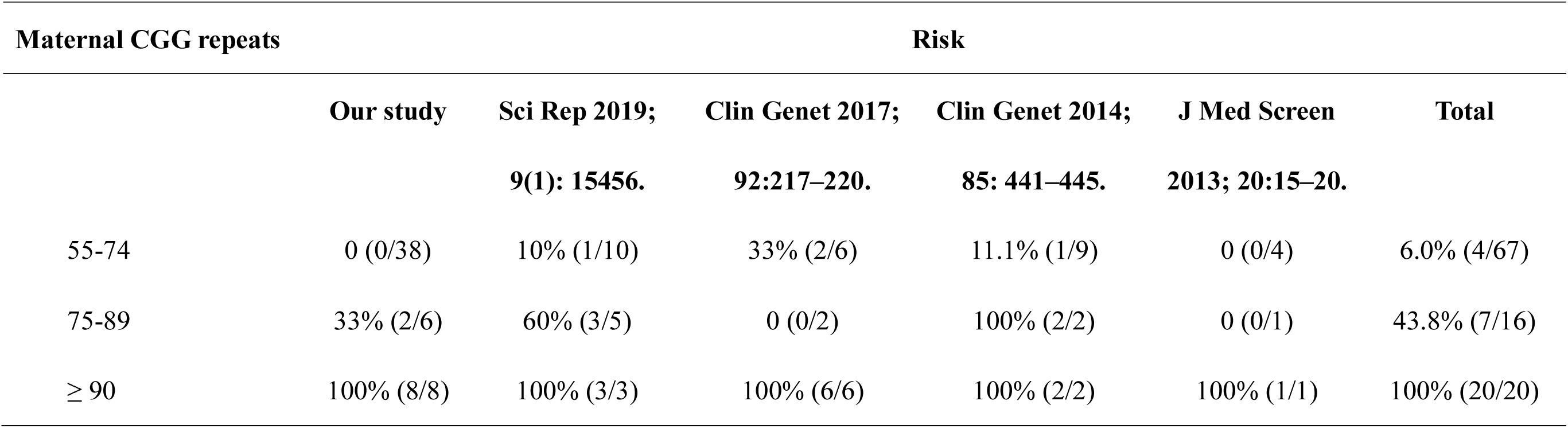
Risk of full mutation expansion for maternal premutation alleles with different CGG repeat lengths.

**Figure 1.**
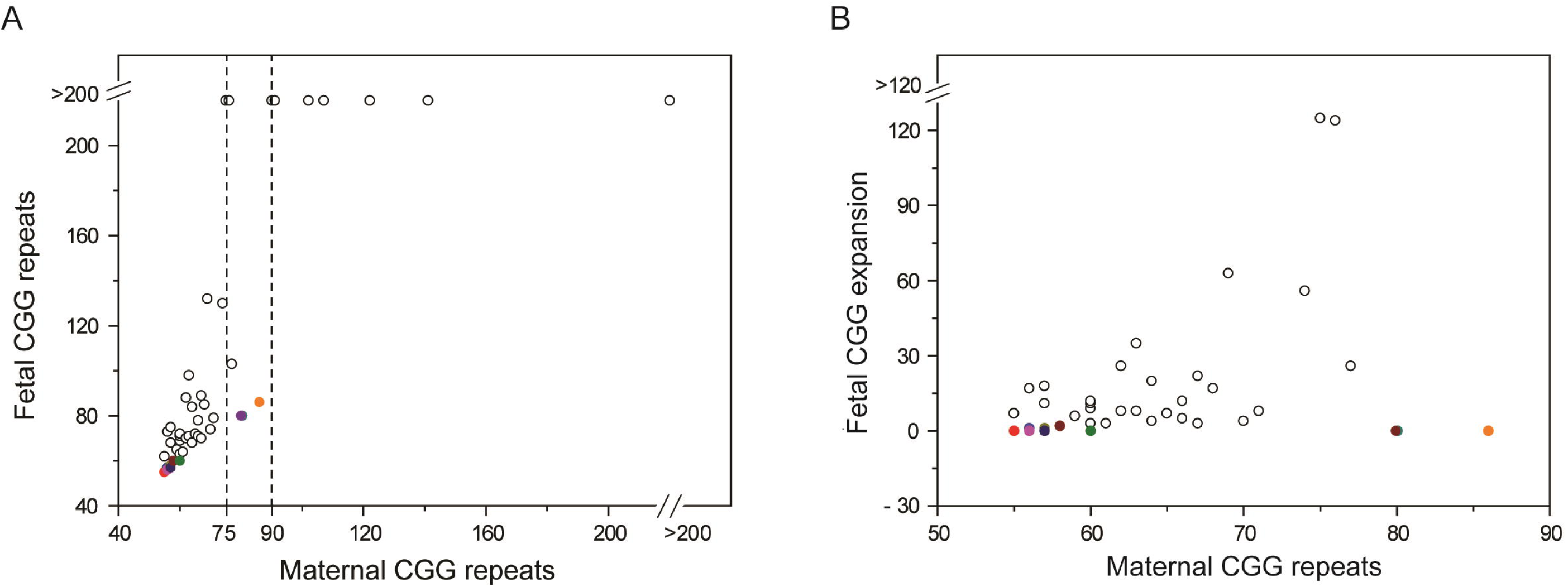
Transmission of intergenerational CGG repeats. (A) Intergenerational CGG repeat transmission profiles of 51 premutation and 2 full mutation alleles. The transmission patterns were uncertain in one mosaic mother and fetus pair. This pair was excluded (Supplemental Table 3). (B) Intergenerational CGG transmission expansion of premutation alleles with CGG repeats under 90. The colored circles denote the alleles with AGG interruptions, while the black circles denote the alleles without AGG interruptions. Alleles ≥200 CGG repeats are the same size to facilitate demonstration.

### Protective effect of AGG interruptions

In cohort 3, AGG interruption was observed in 19 PM alleles out of 87 carriers, of which 13 alleles were transmitted to the 14 fetuses (Supplemental Table 2). Of these 14 transmission events, 10 were stably transmitted, 3 were expanded with 1 CGG copy, and 1 was expanded with 2 CGG copies. In contrast, no stable transmission was observed in the PM alleles without AGG interruptions (Supplemental Tables 2 and 3).

In the subset of maternal PM alleles with 75-89 CGG repeats, 3 PM alleles with AGG interruptions were stably transmitted, while 2 of 3 PM alleles without AGG interruptions expanded to FM alleles (Fig. 1A and Supplemental Tables 2 and 3).

While no AGG interruptions were observed in the PM alleles with CGG repeat sizes ≥90, 100% PM alleles in this subset expanded to FM alleles in intergenerational transmission (Fig. 1A). In terms of PM alleles with CGG repeat sizes under 90, intergenerational CGG expansion was evident in maternal alleles without AGG interruptions compared to their AGG interruption counterparts (Fig. 1B). Based on the data (Fig. 1B), Kruskal–Wallis ANOVA confirmed that the absence of AGG interruptions had a significant e□ect on the risk of intergenerational CGG expansion (P < 0.001).

Lastly, we compared the AGG interruption status between normal, PM, and FM alleles of 105 carriers in cohorts 1 and 2. As shown in Table 6, AGG interruptions were observed in all normal alleles, where the most prevalent number of AGG interruptions was 2. In comparison, the loss of AGG interruption began with the smallest PM alleles (55 CGG repeats), while no AGG interruptions were observed in PM alleles with over 86 CGG repeats. In total, 22 PM alleles had 1 AGG interruption and 1 PM alleles had 2, while no AGG interruptions were observed in the other 82 PM alleles and 4 FM alleles. Chi-square tests confirmed that the absence of AGG interruptions was strongly correlated with PM and FM alleles (P < 0.001).

**Table 6.** Allelic AGG interruption profiles of 105 FXS carriers.

| AGG interruption | Normal allele | PM allele | FM allele<sup>*</sup> |
| --- | --- | --- | --- |
| 0 | 0 | 82 | 4 |
| 1 | 5 | 22 | 0 |
| 2 | 94 | 1 | 0 |
| 3 | 6 | 0 | 0 |

FXS: Fragile X syndrome; PM: pre-mutation; FM: full mutation.
\*Four carriers carried FM alleles, two of which were size mosaics. The CGG repeat sizes of these carriers were 30/>200, 36/>200, 30/141/>200, and 29/84/111/>200, respectively.

### Cost-effectiveness analysis

The decision-analytic model established in this study is shown in Fig. 2. In terms of the population-based screening branch, 71 carriers were identified from 39,458 adult women based on the data of cohort 1. According to the 18.6% incidence of FM fetuses in East Asian populations (Table 4), 13 FM fetuses and 58 normal and PM fetuses were expected from these 71 carriers. In terms of the targeted screening branch, only high-risk women were screened. Based on a previous study, the prevalence of FXS carriers in Taiwanese women with FXS-associated family history was approximately 100 in 725 ^23^. The prevalence of FXS carriers in the East Asian populations without FXS-associated family history was approximately 1/796 (Table 3). Accordingly, approximately 157 women should be high-risk for FXS carriers among 39,458 adult women, with 22 carriers and 4 FM fetuses. Meanwhile, 49 carriers would be missed since the rest of the 39,301 adult women were not tested, which could result in the birth of 9 FM babies. With the assumption of 60% penetrance, 5 babies would be affected with FXS.

**Figure 2.**
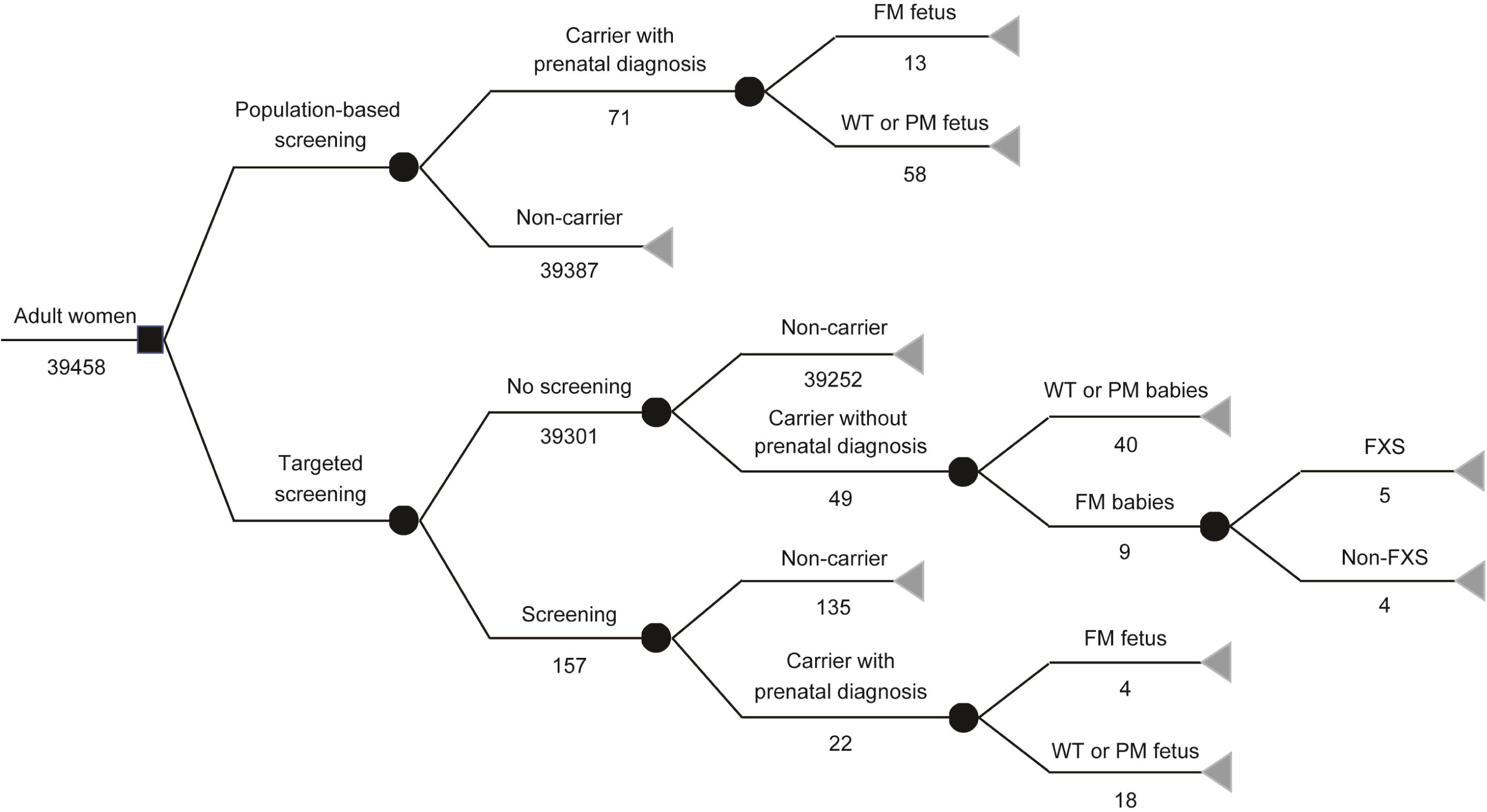
Decision-analytic model for strategies of FXS carrier screening based on the general population and targeted populations. Based on experimental and hypothetical data, the number of subjects is listed.

According to this model, population-based screening and targeted screening strategies to identify a FM fetus would cost approximately 305,707.69 USD and 6125 USD respectively (Table 7). However, the targeted screening strategy would miss approximately 69.2% (9/13) of FM fetuses, suggesting that 1.25 equivalent FXS patients would be born for every FM fetus identified (Fig. 2). Therefore, a potential cost of 625,000 USD would be required due to the life-cycle burden of FXS patients. In this context, population-based screening would be more cost-saving than targeted screening for preventing the birth of FXS patients (Tables 7 and 8). The primary cost in population-based screening is derived from the cost of screening testing, and derived from the life-cycle burden of FXS patients in targeted screening (Table 8). In view of the 100% effectiveness and advanced cost-effectiveness, population-based screening represented a dominant strategy compared to targeted screening.

**Table 7.** Estimated cost for the identification of an FM fetus.

| Screening<br>strategy | Cost | Identified FM<br>fetuses | Cost /<br>identified FM<br>fetuses | Potential<br>cost | Potential cost /<br>identified FM<br>fetuses | (Cost + potential<br>cost) / identified FM<br>fetuses |
| --- | --- | --- | --- | --- | --- | --- |
| Population | 3,974,200 | 13 | 305,707.69 | 0 | 0 | 305,707.69 |
| Targeted | 24,500 | 4 | 6,125 | 2,500,000 | 625,000 | 631,125 |

**Table 8.** Cost-effectiveness of FXS carrier screening.

| Screening strategy | Cost per woman | Potential cost per woman | Effectiveness | Cost per woman / effectiveness | (Cost + potential cost) per woman / effectiveness |
| --- | --- | --- | --- | --- | --- |
| Population | 100.72 | 0 | 1 | 100.72 | 100.72 |
| Targeted | 0.62 | 63.36 | 0.308 | 2.01 | 207.72 |

## Discussion

Understanding the genetic profiles associated FXS is fundamental for optimizing detection tests, genetic counselling, and the intervention strategies in specific populations, especially in East Asian populations, which comprise more than 20% of the global population. As the largest study of carrier screening and prenatal diagnosis for FXS in the East Asian populations to date, our study revealed a 1/556 prevalence of FXS carriers in the general population and a 11.0% incidence of FM fetuses in carrier pregnancies. Several studies have previously reported various levels of FXS carrier prevalence in East Asian populations based on different subject groups and sample sizes. According to these studies, which had relatively large sample sizes (over 5000), the prevalence of FXS carriers in the general population is 1/530-1/788 in South Korea and 1/564 in Northern China ^18, 20, 21^, while the prevalence of FXS carriers in a population without a family history of FXS-associated conditions is 1/781-1/846 in South Korea and 1/776 in Taiwan ^19-21^. Ma et al. reported a higher prevalence (1/410) of FXS carriers in reproductive women in Northern China ^24^. However, the cohort included more than 60% women with a history of spontaneous abortion or induced abortion due to delayed embryo growth. When excluding this subset of samples, the prevalence was adjusted to 1/756. In terms of carrier prevalence in the general population, the differences were not statistically significant among different studies (Table 2; Chi-square test, *P* = 0.690), suggesting a genetic consistency among East Asian populations. Combining this data, the estimated prevalence of FXS carriers is approximately 1/581 in the general East Asian population (Table 2), which is lower than that in Jews and Arabs in Israel ^25^, and pan-ethnic populations in Australia and the USA ^26, 27^, but higher than previous found in the same population derived from relatively small sample sizes ^23, 28^. Noticeably, the difference of carrier prevalence between the general population and populations without a family history is also statistically insignificant (1/581 vs 1/796; Chi-square test, *P* = 0.073), suggesting that the majority of carriers, at least to their knowledge, do not have a family history of FXS-associated conditions, thus minimizing the effectiveness of family history-based screening. Similarly, using the combined data, an estimated 18.6% incidence of FM fetuses in carrier pregnancies (Table 4) and an estimated 1/3124 prevalence of FM fetuses in general pregnancies was suggested, emphasizing the importance of FXS carrier screening.

In prenatal diagnosis, the PM and FM fetuses outnumbered the normal fetuses in our study (Table 1). The combined data (Table 4), which was also indicated in another population-based study ^27^, suggested that compared to normal alleles, PM alleles may be more readily transmitted to the offspring. Expanded *FMR1* alleles have been suggested to be associated with miscarriage ^24, 29^. Our finding supported the concept that miscarriages would more likely be attributed to a disrupted maternal environment rather than developmental defects caused by fetal expanded alleles ^24, 29^. However, FM alleles could be the cause of miscarriage in male fetuses. In our study, female was the predominant gender found in FM fetuses (Table 1), which agreed with a previous study ^19^. Based on the important roles of *FMR1* in fetal development, and the higher penetrance of *FMR1* in males than females, we speculate that male FM fetuses are more readily aborted. However, more data will be required to confirm this speculation. The current categorization of *FMR1* alleles is used worldwide ^5, 15^. However, the thresholds between allele forms may change with increasing empirical data and research, especially in different populations ^5^. In our study, the smallest repeat size of PM alleles that expanded to FM alleles was 75. However, smaller PM sizes associated with FM expansion have been reported in previous studies based on the same population, in which the smallest repeat size was 56 ^19, 20, 23^. In this regard, the currently used cut-off between normal and PM alleles is valid in East Asian population. It is widely known that the number of CGG repeats is an important determinant for assessing the risk of FM expansion in carriers. According to the data, the integral risk of FM expansion is 6.0%, 43.8%, and 100% for PM alleles with 55-74, 75-89, and ≥90 CGG repeats, respectively, in preliminarily assessments (Table 5). Further studies with more cases of CGG intergenerational transmission will be needed to refine the CGG repeat-associated risk assessment further.

As the most promising protective factor, AGG interruption has been widely confirmed to play an important role in CGG transmission stability ^6, 10, 11^. More recently, it was incorporated into the risk assessment of FXS carriers in a population-based screening clinical practice ^26^. However, the protective characteristics of AGG interruption showed differences between Israeli and USA populations ^11^. Moreover, Manor et al. showed that AGG interruptions were more likely a polymorphism rather than a protective factor in individuals of Bedouin ethnicity in Israel ^14^. Similar to carrier prevalence, the role of AGG interruption could be ethnic-specific and will need to be comprehensively evaluated before it can be used for the clinical risk assessment of specific populations ^13^. Unfortunately, studies on the role of AGG interruption in East Asian populations are limited, and AGG interruption is not included in standard genetic counseling due to a lack of validation. In our study, the absence of AGG interruptions had a significant e□ect on the risk of intergenerational CGG expansion (Fig. 1B), and was strongly correlated with the PM and FM alleles (Table 6). Therefore, we have demonstrated that the protective effect of AGG interruption is relevant to the East Asian populations. In particular, AGG interruption seems to play an important role in protecting PM alleles from FM expansion in the subset of maternal PM alleles with 75-89 CGG repeats (Fig. 1A; statistical analysis was not performed due to the limited sample size). Moreover, the percentage of PM alleles without AGG interruptions (3/6) was comparable to the integral risk of FM expansion (43.8%) in this subset (Table 5). These results further supported that AGG interruption may play an important protective role in this PM range. However, future studies with more cases of AGG interruption with CGG transmissions will be needed to refine the AGG interruption-associated risk assessment further. Although the benefits of the prenatal intervention of FXS are widely acknowledged, population-based FXS carrier screening has yet to be officially endorsed by professional guidelines. This strategy lacks a consensus with regards to some of the main concerns, including lack of disease education, lack of cost-benefit or cost-effectiveness advantages, difficulties in genetic counseling, and potential for psychosocial harms ^30, 31^. Instead, targeted screening based on the family history of FXS-association conditions is currently recommended ^15^. As mentioned above, the insignificant difference of carrier prevalence between the general population and populations without a family history of FXS suggested that family history is not an effective indicator for FXS carrier screening in East Asian populations. In our model, we calculated that targeted screening misses approximately 69% of carriers (Fig. 2). Moreover, our findings were in accordance with those reported by a study on an Australian population ^27^. Based on the genetic profiles of the East Asian population, we confirmed that population-based screening is more cost-effective than targeted screening (Tables 7 and 8). The primary cost of population-based screening is the cost of screening testing (Table 8). The cost of genetic testing had tended to decrease with rapid developments in biotechnologies, indicating that a greater economic advantage could be expected in the near future. Furthermore, the development of novel screening methods, such as “next- and third-generation sequencing” ^32, 33^, may also improve the testing capacity, thus further advancing the implementation of population-based screening to large populations. Several issues that contribute to the cost of population-based screening, including the infrastructure and human resources needed to provide appropriate education, counselling, interventions, and follow-up, were not included in our simplified model. However, the unit cost would be limited in FXS screening due to the fact that investment is not specific for a single condition but benefits numerous genetic diseases. This is particularly pertinent in the era of expanded population-based carrier screening, where hundreds of diseases can be screened simultaneously ^26^. In addition to its cost-effectiveness, other factors point towards the expansion of FXS screening criteria ^34, 35^. In fact, several national population-based screening programs, including FXS carrier screening, have already been implemented and have obtained positive outcomes ^25-27^. The development of an effective FXS treatment could drastically improve the health of patients and lower disease-associated expense, and could revise the strategies for FXS intervention. Nonetheless, in the current stage, population-based carrier screening is the dominant strategy for FXS intervention in East Asian populations.

In conclusion, we reported on the largest carrier screening and prenatal diagnosis for FXS in East Asian populations to date, delineating the FXS-associated genetic profiles in this population. Our findings provide evidence that supports the implementation of population-based carrier screening over family history-based targeted screening for FXS intervention.

## Acknowledgments

We thank all the participants for their kind support of this work.

